# Neural effects of placebo analgesia in fibromyalgia patients and healthy individuals

**DOI:** 10.1101/2020.07.31.231191

**Authors:** Eleni Frangos, Marta Ceko, Binquan Wang, Emily A. Richards, John L. Gracely, Luana Colloca, Petra Schweinhardt, M. Catherine Bushnell

## Abstract

Placebo analgesia is hypothesized to involve top-down engagement of prefrontal regions that access endogenous pain inhibiting opioid pathways. Fibromyalgia (FM) patients have neuroanatomical and neurochemical alterations in pathways relevant to placebo analgesia. Thus, it remains unclear whether placebo analgesic mechanisms would differ in FM patients compared to healthy controls (HCs). Here, using placebo-analgesia-inducing paradigms that included verbal suggestions and conditioning manipulations, we examined whether behavioral and neural placebo analgesic responses differed between 32 FM patients and 46 age- and sex-matched HCs. Participants underwent a manipulation scan, where noxious high and low heat were paired with the control and placebo cream, respectively, and a placebo experimental scan with equal noxious heat temperatures. Before the experimental scan, each participant received saline or naloxone, an opioid receptor antagonist. Across all participants, the placebo condition decreased pain intensity and unpleasantness ratings, decreased activity within the right insula and bilateral secondary somatosensory cortex, and modulated the Neurologic Pain Signature. There were no differences between HCs and FM patients in pain intensity ratings or neural responses during the placebo condition. Despite the perceptual and neural effects of the placebo manipulation, prefrontal circuitry was not activated during the expectation period and the placebo analgesia was unaltered by naloxone, suggesting placebo effects were driven more by conditioning than expectation. Together, these findings suggest that placebo analgesia can occur in both HCs and chronic pain FM patients, without the involvement of opiodergic prefrontal modulatory networks.

## INTRODUCTION

Multiple lines of evidence suggest that placebo analgesia has a neurobiological basis involving the engagement of endogenous pain inhibitory systems that likely dampen afferent nociceptive input [1; 16; 17; 46]. Experimental studies of placebo analgesia report activations during placebo-induced expectation of pain relief throughout regions involved in descending modulatory control of afferent nociceptive input, including the dorsal lateral and ventral medial prefrontal cortex (DLPFC, VMPFC) [19; 43]. Consistent with this idea, transient inhibition of the DLPFC using repetitive transcranial magnetic stimulation has been shown to block placebo analgesia [26], and the degeneration of the prefrontal lobes that occurs in Alzheimer’s disease abolishes placebo analgesic effects [7]. Other studies show the endogenous release of opioids during placebo analgesia [46; 49], suggesting a possible neurochemical mechanism. The role of endogenous opioids in placebo analgesia created through enhancing expectations has been directly confirmed by showing that naloxone, an opioid receptor antagonist, can block expectation-related placebo effects [1; 16; 29]. In contrast, it seems that placebo analgesia created primarily through conditioning does not involve the opioid system as long the unconditioned stimulus is not an opioid [5].

Despite this evidence, the prevailing idea that placebo analgesia involves activation of endogenous descending-control systems and subsequent dampening of afferent nociceptive pathways has recently been challenged by a meta-analysis that revealed inconsistent effects within the neurologic pain signature (NPS) network. The authors propose that placebo treatments affect pain via brain mechanisms largely independent of effects on bottom-up nociceptive processing [50]. Nevertheless, given the small sample size of many studies in the meta-analysis, differences in placebo induction methods and pain-evoking stimuli, and the large network comprising the NPS, the effects of placebo analgesia on bottom-up nociceptive processing is still unclear.

Another unresolved question involves possible differences in placebo analgesia mechanisms between chronic pain patients and healthy individuals. A recent meta-analysis concluded that pain patients have greater benefit from placebo treatment than healthy individuals and that placebo studies on only healthy individuals may underestimate the magnitude of the placebo analgesic effect [18]. This finding is intriguing since evidence shows that chronic pain patients have anatomical, functional and neurochemical alterations in brain regions involved in placebo analgesia [3; 10; 21; 22; 27; 31; 36; 47]. Furthermore, expectation levels may be altered in patients because of their experience with pain and effectiveness of medications, which could either increase or decrease placebo effectiveness [13]. To date, very few studies have directly compared placebo effects in chronic pain patients to healthy people using the same experimental paradigm, and only half of the studies examined neural effects [12; 24; 28; 30].

Thus, the current study investigates perceptual effects and neural mechanisms of placebo analgesia in chronic pain patients diagnosed with fibromyalgia (FM) and matched healthy controls (HCs). We asked the following questions: Q1) Does the magnitude of perceptual placebo analgesia differ between HCs and FM patients? Q2) Does placebo analgesia involve different brain regions for HCs and FM patients? Q3) Does opioid-receptor blockade differentially affect (conditioned) placebo analgesia in HCs and FM patients?

## METHODS

### Participants

Based on the sample size calculation described below for the main effect of cream on perceptual placebo effects (control cream vs. placebo cream), we enrolled a minimum of 40 participants per group. A total of 96 participants were enrolled in the study. Eighteen participants were not included due to attrition, technical problems resulting in incomplete experimental sessions, or because they did not meet the inclusion criteria listed below during a second screening (e.g., pain intensity rating on the day of fMRI scanning of >4 out of 10 for the patients). Thus, the final study population included 32 chronic pain patients diagnosed with FM (30 females, 2 males, mean age ± SD: 43 ± 12.3 years, range 24-62 years) and 46 age- and sex-matched healthy controls (39 females, 7 males, 40 ± 13 years, range 19-64; p=0.34). Groups were further matched on race, level of education, and level of physical activity (International Physical Activity Questionnaire IPAQ [8] (Table 1).

**Table 1.**
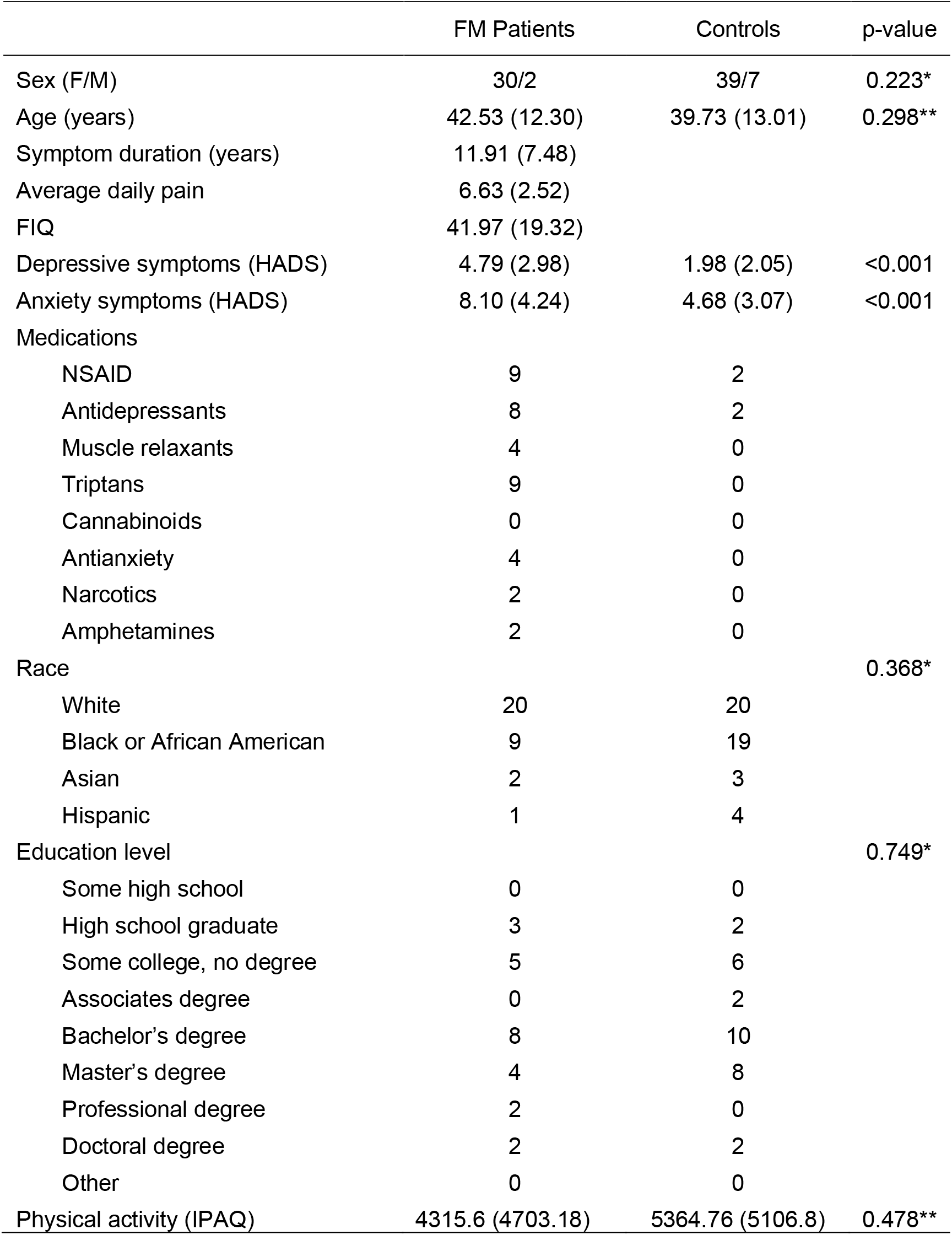
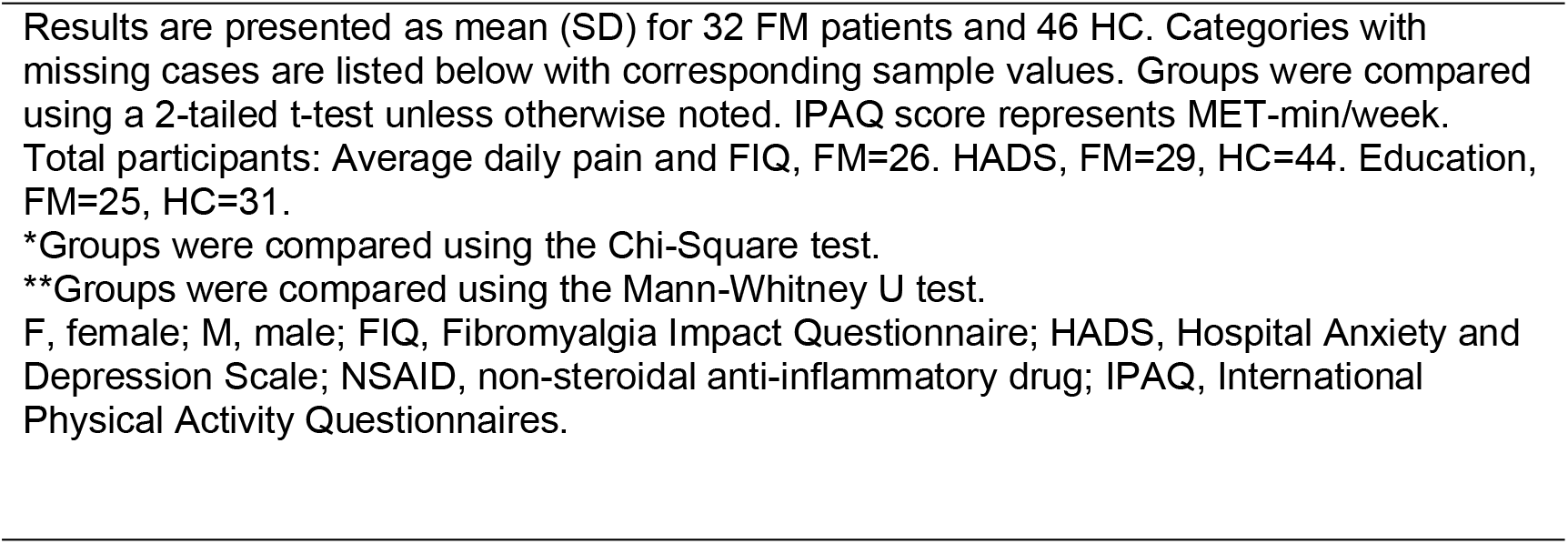
Demographic and clinical data for fibromyalgia (FM) patients and controls.

|  | FM Patients | Controls | p-value |
| --- | --- | --- | --- |
| Sex (F/M) | 30/2 | 39/7 | 0.223* |
| Age (years) | 42.53 (12.30) | 39.73 (13.01) | 0.298** |
| Symptom duration (years) | 11.91 (7.48) |  |  |
| Average daily pain | 6.63 (2.52) |  |  |
| FIQ | 41.97 (19.32) |  |  |
| Depressive symptoms (HADS) | 4.79 (2.98) | 1.98 (2.05) | <0.001 |
| Anxiety symptoms (HADS) | 8.10 (4.24) | 4.68 (3.07) | <0.001 |
| Medications |  |  |  |
| NSAID | 9 | 2 |  |
| Antidepressants | 8 | 2 |  |
| Muscle relaxants | 4 | 0 |  |
| Tryptans | 9 | 0 |  |
| Cannabinoids | 0 | 0 |  |
| Antianxiety | 4 | 0 |  |
| Narcotics | 2 | 0 |  |
| Amphetamines | 2 | 0 |  |
| Race |  |  | 0.368* |
| White | 20 | 20 |  |
| Black or African American | 9 | 19 |  |
| Asian | 2 | 3 |  |
| Hispanic | 1 | 4 |  |
| Education level |  |  | 0.749* |
| Some high school | 0 | 0 |  |
| High school graduate | 3 | 2 |  |
| Some college, no degree | 5 | 6 |  |
| Associates degree | 0 | 2 |  |
| Bachelor's degree | 8 | 10 |  |
| Master's degree | 4 | 8 |  |
| Professional degree | 2 | 0 |  |
| Doctoral degree | 2 | 2 |  |
| Other | 0 | 0 |  |
| Physical activity (IPAQ) | 4315.6 (4703.18) | 5364.76 (5106.8) | 0.478** |

Results are presented as mean (SD) for 32 FM patients and 46 HC. Categories with missing cases are listed below with corresponding sample values. Groups were compared using a 2-tailed t-test unless otherwise noted. IPAQ score represents MET-min/week. Total participants: Average daily pain and FIQ, FM=26. HADS, FM=29, HC=44. Education, FM=25, HC=31.
\*Groups were compared using the Chi-Square test.
\*\*Groups were compared using the Mann-Whitney U test.
F, female; M, male; FIQ, Fibromyalgia Impact Questionnaire; HADS, Hospital Anxiety and Depression Scale; NSAID, non-steroidal anti-inflammatory drug; IPAQ, International Physical Activity Questionnaires.

The inclusion criteria for patients included a diagnosis of FM (excluding other pain disorders) confirmed by medical records or directly by the treating physician, and chronic widespread pain for at least 1 year with an average daily pain intensity of at least 4 out of 10. Exclusion criteria for all participants included smoking of >10 cigarettes/week, alcohol consumption of >7 drinks/week for women and >14 drinks/week for men, use of recreational drugs, use of opioid medication, pregnancy or breastfeeding, allergies to skin creams and lotions, chronic pain conditions (other than FM for patients), major medical, neurological, or current psychiatric conditions, including severe depression and generalized anxiety disorder, and MRI contraindications. Additional exclusion criteria for healthy controls included taking any pain medication other than NSAIDs within the last month or for more than one month on a continual basis within the last 6 months. Patients remained on their regular pain medications (Table 1).

The study received approval from the NIH Institutional Review Board (IRB), and written informed consent was obtained from all participants according to the Declaration of Helsinki. As per IRB guidelines, the consent form included a general statement about deception: “At some point during the study we will give you misleading information. After the study is finished and all participants have been tested, we will explain how the information was not true and why.” No further details regarding deception were provided, and participants were not informed that the purpose of the study was to investigate placebo analgesia. Participants were compensated for completion of the study.

#### Sample size calculation

Our primary outcome measure was the perceptual effect of placebo analgesia as measured by pain intensity ratings. Thus, an *a priori* sample size calculation was conducted, based on the results conducted by our group on healthy participants [35]. In order to detect a 10-point difference with a standard deviation of 22 on the 100-point VAS between the placebo cream and hydrating cream, fixing the statistical power to 80% and the Type I error probability to α=0.05, we determined that 40 participants would be needed to detect a placebo effect. The following formula was used for the sample size estimation for within-subject comparisons: n= 2 + C (s/d)2, where C is a constant of 7.85 when α=0.05 and β=0.80, s represents the standard deviation of the individual measurements, and d represents the expected mean difference to be detected.

### Study design

This study was patterned after a well-established placebo analgesia paradigm including both expectation and conditioning components in a between- and within-participants design [13; 16; 45], but participants were given additional instructions to create more neutral expectations. Participants were told that the study was designed to investigate the effect of an opiate antagonist, naloxone, on the pain-reducing effects of a powerful topical local analgesic cream (“NIH-compound”, in reality, the placebo cream), as well as on brain responses to painful and non-painful cutaneous thermal stimuli in chronic pain patients and healthy control participants. They were told (truthfully) that the study was double blind and that they would have a 50% chance of receiving naloxone. They were further told that naloxone may or may not block the “analgesic” effect of the cream. These instructions are in line with instructions received in clinical trials of drug efficacy but contrast those of most other experimental placebo studies in which participants are told that they are receiving a “powerful analgesic”, without any additional stipulations, e.g., the possibility that the naloxone might reverse the “analgesic” effects of the cream, as in the present study [13; 16; 45]. In addition, in order to minimize expectations that patients may have had about the applicability of the analgesic cream to their clinical pain states, we indicated that the analgesic cream was for applications to small skin areas and thus not suitable for the more widespread pain of deeper structures characteristic of FM.

The experimental design and paradigm are described below and detailed in Figure 1. The experiment took place across two separate days and included the following: Day 1-medical exam and questionnaires; in the mock scanner-pain calibration and placebo manipulation 1; Day 2-in an MRI scanner-a conditioning scan (placebo manipulation 2), drug administration, a high resolution anatomical scan, and two placebo test scans. Thermal stimuli (4-8.5 seconds) were applied to four different 4×4 cm placebo cream or control cream treated areas of the lower left leg (Fig. 1) using a contact thermode (ATS, Medoc Pathway Model, Medoc Ltd Advanced Medical System, Israel) on both days. Participants rated pain intensity and unpleasantness each on a visual analog scale (VAS) previously validated to be sensitive to subtle psychological manipulations [41; 42] (anchors: pain intensity, 0=no sensation, 100=pain threshold, 200=intolerable pain; unpleasantness/pleasantness, −100=extremely unpleasant, 0=neutral, 100=extremely pleasant).

**Figure 1.**
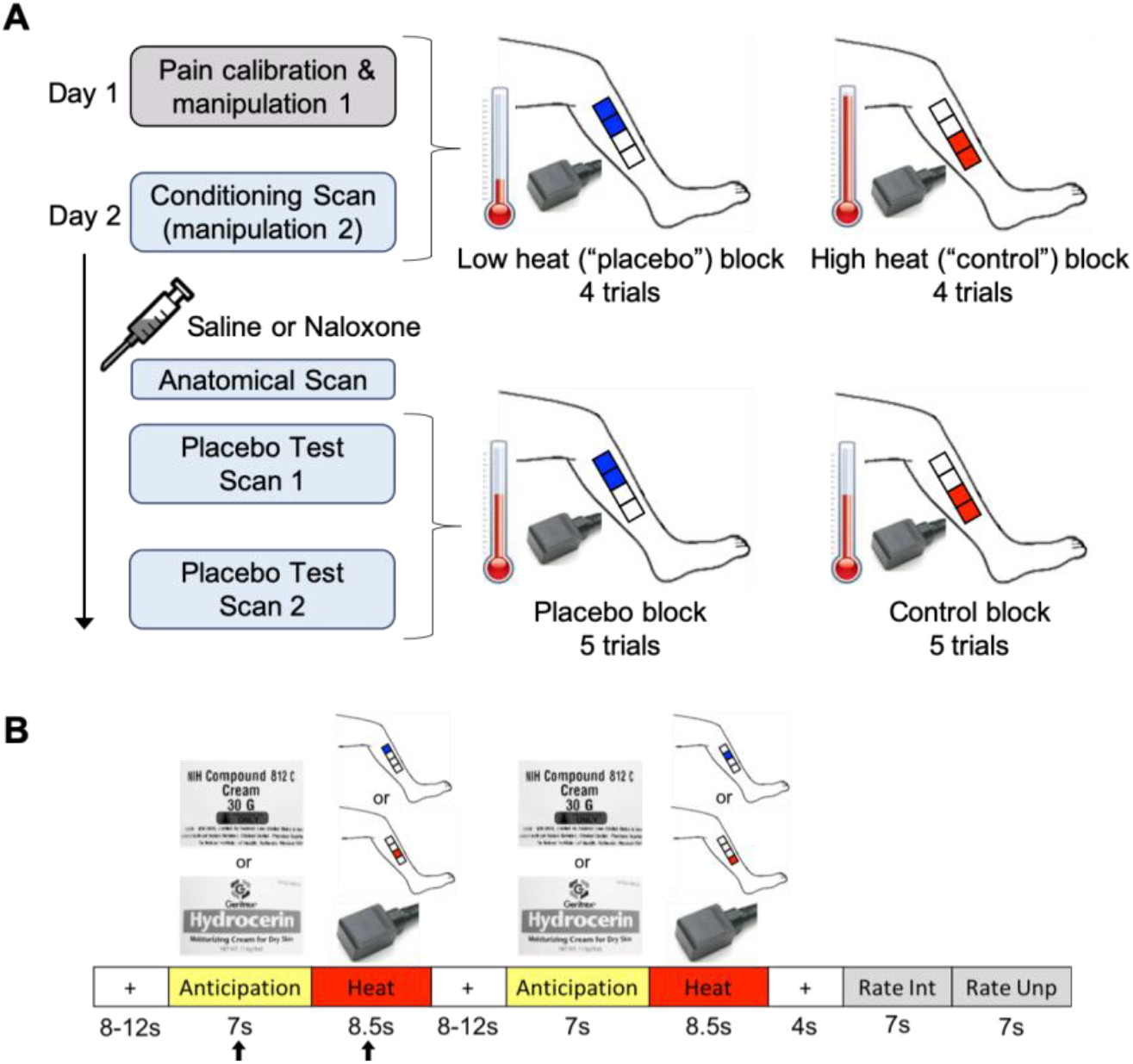
Experimental design and paradigm. A. Day 1 included a pain calibration procedure to determine which temperatures participants would rate within the low pain range and high pain range on a 0-200 VAS pain intensity scale (0=no sensation, 100=pain threshold, 200=intolerable pain). The low and high temperatures, differing by ~2°C for each individual, were used during manipulation sessions 1 and 2. Manipulation session 1 was conducted in a mock scanner. Participants were presented with the *powerful topical local analgesic* “NIH-compound” (placebo) cream and the “hydrating” (control) cream (both creams were identical and inert). Each cream was applied to two 4×4cm areas of the lower left leg (distal vs. proximal placement was randomized between participants) and was left on for 5 min before wiping off. To condition and convince participants of the effectiveness of the “analgesic”, the individualized low and high heat stimuli were applied to the placebo and control cream locations, respectively (4 trials per condition, two mock scan runs). One run of the manipulation procedure took place on Day 2 in an fMRI scanner followed by a bolus of saline or naloxone, an anatomical scan, and two placebo test scans. The heat temperature administered during the placebo test scans was mid-way between the individualized low and high temperatures that were used during the manipulation session, and most importantly, the *same* temperature was used for the placebo and control condition. Each condition block had 5 trials per scan. Participants were pseudo-randomly assigned to receive either the placebo or control block first. B. The trial paradigm during the manipulation and placebo test sessions consisted of a jittered inter-stimulus interval (ISI) of a black crosshair on white background, an anticipation period preceding the heat pulse (cue of either the “NIH-compound” (placebo) cream label (top image) or the “hydrating” (control) cream label (bottom image)), a heat pulse on the left leg during which a thermode image was shown, a second jittered ISI, a second anticipation cue and heat pulse, a post-stimulus ISI, and a rating scale for pain intensity and pain unpleasantness. The black arrows indicate the anticipation period and heat pulse reported in the results.

#### Day 1 - mock scanner

##### Pain calibration procedure

A sequence of brief heat stimuli (4-7seconds) between 37° and 50°C was presented to four 4×4cm calibration areas on the right leg. After each stimulus, participants rated the pain intensity on the VAS. The data from this phase were used to determine, by interpolation, temperatures that the subject would rate as low-pain (VAS ~110-130) and high-pain (VAS ~150-180). These individually calibrated temperature intensities were used during the manipulation phases (top half of Fig. 1A).

##### Placebo manipulation 1

The placebo manipulation procedure (top half of Fig. 1A) mainly consisted of conditioning through surreptitious temperature manipulation. In contrast, we only induced limited expectations of pain relief through instructions that participants may or may not receive naloxone which may or may not block the analgesia. The “NIH-compound” (placebo) cream was applied to two of the four 4×4 cm marked regions on the lower left leg and the “hydrating” (control) cream, used “to control for moisturizing effects”, was applied to the other two marked regions (Fig. 1A). In actuality, the two creams were the identical moisturizing cream. The distal vs. proximal placement of the creams was randomized between participants. The creams were left on the skin for 5 minutes, with the instruction that this time insured full absorption of the “analgesic” cream. The creams were then carefully removed from the skin, and the conditioning manipulation phase started. During this phase, participants were lying in a supine position in the mock scanner (MRI Simulator, Psychology Software Tools) to allow for habituation to the scanning procedure, with eyes focused on a computer screen while undergoing the heat stimulation paradigm described below and in Figure 1B. Unbeknownst to participants, the temperature presented on the “analgesic” region was lower (VAS ~110-130) compared to the temperature presented on the “hydrated” region (VAS ~150-180). This served to reinforce the verbal suggestion that the placebo cream was an effective analgesic cream and to create conditioning independent of suggestions, which is also known to produce placebo analgesia without explicit manipulations of expectations of pain relief [1; 5; 23]. Two runs were conducted in the mock scanner. Each run had 4 trials per condition (trial paradigm detailed below).

##### Trial paradigm

Each trial (Fig. 1B) consisted of a baseline period (jittered 8-12 sec; black crosshair on white background), an anticipation period (7 sec; grayscale picture of control cream or placebo “analgesic” cream), a heat pulse (8.5 sec; gray scale picture of thermode), a second anticipation period and heat pulse, a post-stimulus rest period (4 sec; black crosshair on white background), and two rating periods (7 sec for intensity, 7 sec for unpleasantness; black VAS on white background). Each heat pulse was presented on one of two pairs of treated 4×4 cm regions of the lower left leg.

#### Day 2 - fMRI scans

##### Placebo manipulation 2

On Day 2, participants underwent a second placebo manipulation procedure (conditioning scan) to further reinforce conditioning. The drug administration, high resolution anatomical scan and placebo test phase shortly followed. Just prior to the conditioning scan, four 4×4cm areas were marked on the left leg (Fig.1A) and an intravenous line was inserted into the right arm. As on Day 1, the two creams were applied in the same manner, and temperature presented for the placebo cream condition was lower compared to the temperature presented for the control cream condition. There were 4 trials per condition following the trial paradigm described above (Fig. 1B). Next, participants were given either a bolus injection of saline or naloxone, with approximately half of the participants in each group (HC or FM) randomized to receive naloxone and the other half saline. An anatomical MRI was acquired while the drug reached its peak effect.

##### Placebo test phase

The first scan of the placebo test phase then began. Each condition had 5 trials (Fig.1B) but the heat stimuli were now the same temperature for both conditions, i.e., a painfully hot temperature mid-way between the individualized low and high temperatures administered during the manipulation phase, following the design of Eippert et al. [16]. Half of the participants from each group were pseudo-randomly assigned to receive the first block of 5 trials with stimuli presented on the placebo cream site and the other half on the control cream site. A second placebo test scan followed immediately after the first, with the order of cream sites reversed for each participant.

##### Drug administration

Approximately 10 min prior to the placebo test scan, some participants (23 HCs and 20 FM patients) received an infusion of naloxone, and the others (23 HCs and 12 FM patients) received an infusion of saline. Naloxone or saline was administered by the NIH Clinical Center nursing staff in a double-blinded fashion using block-stratified (age, sex) randomization. Participants were informed about naloxone, including its pharmacological properties, general clinical use, and possible side effects. Participants were also informed that they would most likely not notice that they had received naloxone as it generally has no noticeable effects in the dose employed. To achieve a constant plasma level throughout the ~40 min testing phase, a bolus dose of naloxone (0.05 mg/kg bodyweight; generic) or saline was first administered via an intravenous line, followed by an intravenous infusion dose of 0.08 mg/kg/hr naloxone or saline (diluted in 250 ml of saline), starting immediately after the bolus injection and continuing for ~40 min. The total dose of naloxone could not exceed 10 mg, a dosage used clinically to reverse the effects of opiates in opiate-overdose (Micromedex 2), as required by NIH IRB guidelines. Vital signs (blood pressure, pulse, respiration, and pulse oximetry) were taken once before the intravenous bolus, once 5 minutes thereafter, and once post infusion. Participants were asked to rate naloxone-related adverse effects (dry mouth, dry skin, blurred vision, sedation, nausea, dizziness, headache) on a scale from 0=non-existent to 6=extremely strong [16]. The drug administration procedure was similar to that used to block placebo analgesia in young male healthy volunteers [16], except that the dose of naloxone was lower in the present study (present study vs. Eippert et al. [16], bolus dose: 0.05 mg/kg vs. 0.15 mg/kg; intravenous infusion dose: 0.08 mg/kg/hr vs 0.2 mg/kg/hr), as required by NIH IRB guidelines. Both naloxone and saline were well tolerated, and the expected side effects were non-existent to minimal for the two drugs across all participants (Supplementary Table 1). No significant adverse events were observed.

##### Randomization and allocation

Both the saline and naloxone solutions were prepared by the NIH pharmacy and furnished in individual subject doses on the day of the experiment for each subject. The NIH pharmacy provided subject randomization and maintained the randomization code until completion of data collection.

##### Effectiveness of cream and desire for pain relief

Numerical rating scales were used to assess the perceived effectiveness of the “NIH analgesic” (anchors: 0=not effective at all, 10=the most effective) and the desire for pain relief during the heat stimulation (anchors: 0=no desire for pain relief, 10=the most intense desire for pain relief imaginable). These scales were administered after manipulation 1, after manipulation 2, and after the placebo test scans.

##### Current pain and discomfort

To assess possible interference of ongoing pain and discomfort with experimental pain ratings and fMRI findings, the intensity of current pain and discomfort during the fMRI data acquisition were assessed once immediately following the manipulation fMRI scan and once immediately following the two runs of the placebo-test fMRI scan. An 11-point numerical intensity rating scale from 0-10 was used for both pain (0=no pain, 1=pain threshold, 10=worst bearable pain) and discomfort (0=no discomfort, 1=discomfort threshold, 10=worst bearable discomfort). In addition, type and location of pain and of discomfort were assessed. These scales were administered after placebo manipulation 1, after placebo manipulation 2, and after the placebo test scans.

### fMRI acquisition

All participants completed a 9.5 min fMRI scan during the placebo conditioning phase, a 4.5-min high-resolution anatomical MRI scan, and two 11.8 min fMRI scans during the placebo test phase. Throughout the session, participants wore earplugs and their heads were immobilized. Brain images were acquired using a 3 Tesla Siemens Skyra MRI scanner (Siemens, Erlangen, Germany) with a 20-channel head and neck coil. Structural MRI images (T1-weighted) were acquired using an MPRAGE sequence (TR = 1900 ms, TE = 2.07 ms, flip angle = 9°, 1mm isotropic voxels, image matrix = 256×256, 192 slices). Functional MRI data were acquired using a blood oxygenation level-dependent (BOLD) protocol with a T2*-weighted gradient echo planar imaging (EPI) sequence (TR = 2000 ms, TE = 29 ms, flip angle 70°, 3.5 mm isotropic voxels, FOV = 64×64, 38 slices). Axial slices were oriented 30 degrees from the line between the anterior and posterior commissures, covering the entire brain, and excluding the eyes. After discarding the first three volumes to allow for steady-state magnetization, 285 volumes and 355 volumes were acquired for the conditioning and placebo test scans, respectively. During the fMRI scans, heart rate, blood oxygenation and respiration were monitored.

### Behavioral data analysis

Outcome measures were compared using independent samples two-tailed t-tests or a repeated measures analysis of variance (rm-ANOVA) with one within-subject factor (“cream”: placebo cream vs. control cream) and two between-subject factors (“group”: FM vs. HC; “drug”: saline vs. naloxone) in SPSS 25 (IBM). Mann Whitney U tests were used for non-normally distributed data and Chi-square tests were used for categorical comparisons. Correlations between behavioral measures, FM characteristics, and gray matter volume (GMV) were investigated using Pearson correlations. A significance level of p<0.05 was used in all analyses. All results are presented as mean ± SD.

### Voxel-based morphometry (VBM): preprocessing and analysis

In order to evaluate whether the FM patients in the current study had forebrain anatomical changes similar to those reported in other studies [10; 27], VBM analysis was performed. Since brain gray matter volumes are strongly influenced by age [48], we selected a sub-set of the healthy controls that were age- and sex-matched to the patients on an individual basis. Thus, the VBM analysis included 31 FM patients and 31 healthy controls. Structural data were analyzed using FSL-VBM [15; 20] and carried out with FSL tools [37]. Non-brain tissue was removed from the structural images, followed by gray matter segmentation, and non-linear registration to the MNI 152 T1 2mm standard space template [2]. The gray matter images were averaged and flipped along the x-axis to create a left-right symmetric, study-specific gray matter template using 31 FM patients and 31 HC. All native gray matter images were then non-linearly registered to this study-specific template and modulated to correct for local expansion (or contraction) due to the non-linear component of the spatial transformation. The modulated gray matter images were then smoothed with an isotropic Gaussian kernel with a sigma of 4 mm (approx. FWHM = 8 mm). Finally, a voxelwise general linear model (GLM) was applied using permutation-based non-parametric testing, correcting for multiple comparisons across space. A voxel threshold of z > 3.1 and cluster threshold of p < 0.05 was used (results for corrections using z > 2.3 and TFCE are available in the Supplemental material). Age (mean-centered across all participants) was included as a nuisance regressor in the GLM. A 6mm sphere was used to extract the gray matter volume (GMV) values of the DLPFC cluster identified by the results of the contrast HC > FM. Placement of the sphere around the peak voxel (MNI coordinate x = 60, y = 65, z = 58) resulted in overlap with white matter. Therefore, to avoid overlapping the sphere with white-matter, the sphere was moved to MNI coordinate x = 62, y = 65, z = 60, towards the center-of-gravity of the cluster, which included the peak voxel. Based on the a priori hypothesis that FM decreases in gray matter volume would negatively correlate with symptom duration as reported in previous studies [10; 27], a one-tailed partial correlation controlling for age was conducted between FM DLPFC GMV and symptom duration. A significance level of p<0.05 was used.

### fMRI data preprocessing and analysis

All fMRI data were preprocessed and analyzed in FSL (version 5.0.8). Preprocessing included non-brain tissue removal, spatial smoothing (Gaussian kernel of FWHM = 5mm), high pass temporal filtering, six-parameter (3 translations and 3 rotations) rigid body correction for head motion, co-registration to the T1-weighted anatomical image and spatial normalization to the MNI152 T1 2mm template using a 12-parameter linear registration.

Explanatory variables (EVs) were modelled at the individual level using a double-gamma hemodynamic response function. The components within each trial (Fig. 1B) were modelled separately per condition, i.e., the first anticipation period, first heat pulse, second anticipation period, and second heat pulse that occurred within a trial were each modelled as separate EVs for each condition (placebo cream or control cream). The pain rating periods (‘Intensity’, ‘Unpleasantness’) were combined into one EV for each condition (placebo cream or control cream). The fixation periods were not modelled. All anticipation periods and heat periods were included in the model. In total, the model included 10 anticipation periods and heat pulses per condition.

Higher level group analyses were carried out using FSL’s FLAME 1+2 mixed effects modelling to assess group, drug, and group X drug interactions (repeated-measures ANOVA) of the following contrasts for each anticipation period and heat stimulus: placebo > baseline, control > baseline, placebo > control, control > placebo. Pain intensity ratings were included in the model as a covariate of interest for the placebo analyses. For all scans, heat pulse 1 of each trial, regardless of condition, produced significantly greater activity than heat pulse 2 despite movement of the thermode from one 4×4 cm marked area to the second after each heat pulse. In addition, no placebo effects were observed during the second experimental scan and all pain ratings were significantly lower during the second experimental scan compared to the first (see Supplementary section 3). Thus, to account for the habituation effect observed during the second pulse of each trial and the during second scan, only the first anticipation period and first heat stimulus of each trial (black arrows in Fig. 1B) for each condition in the first experimental scan are reported in the main text, with the second pulse and second scan reported in the supplementary material. Specifically, while the model included all 10 anticipation periods and heat pulses, the reported results in the main text are based on heat pulse 1 of each trial, i.e., 5 anticipation periods and 5 heat pulses for each condition of the first scan. Voxel-wise thresholds were set to z > 3.1 or z > 2.3 to assess subtle effects and reduce false negatives (Type II error). All contrasts were cluster-corrected for multiple comparisons across the whole brain at p < 0.05.

### Neurologic Pain Signature

Because of the meta-analysis [50] reporting that placebo analgesia does not affect the NPS, we tested whether the NPS was affected by the placebo cream in our study. To compute the magnitude of the NPS response, the voxel-wise pattern of regression weights was multiplied with each of the following contrast of parameter estimates (“COPEs” from the individual-level analyses) to produce dot products which were then averaged across all voxels resulting in one value for each subject and condition: placebo test scan-control cream > placebo cream, control cream > baseline, placebo cream > baseline. The analyses were completed on MATLAB R2018a (MathWorks) using code provided by the Wager lab (https://canlabweb.colorado.edu/). Differences between the control cream and placebo cream condition were assessed using two-tailed t-tests for dependent samples. A significance level of p < 0.05 was used.

## RESULTS

### Patient characteristics

FM patients had mild to moderate fibromyalgia (FIQ score mean ± SD, 41.97 ± 19.32; range 8-89), and their average symptom duration was 11.91 ± 7.48 years (range 2-30 years) with an average intensity of 6.63 ± 2.52 for daily pain. We found no significant correlations between FM characteristics and placebo effect, i.e., the difference between ratings of the control and placebo creams (see Supplementary Table 2). As expected, the Hospital Anxiety and Depression Scale (HADS) indicated that patients had significantly increased, but sub-clinical, levels of anxiety and depressive symptoms compared to healthy controls. A summary of the clinical and demographic information can be found in Table 1.

To further determine if the FM patients in the current study were similar to FM patients in other studies, we used VBM to determine whether FM patients had previously observed gray matter abnormalities [10; 27]. The whole-brain VBM analysis (corrected for age) performed on 31 FM patients and 31 age- and sex-matched HCs revealed a significant decrease in gray matter volume (GMV) within the left DLPFC of the FM patient group, as indicated by the contrast of HC > FM patients (Fig. 2A). The GMV within the left DLPFC of FM patients was negatively correlated with symptom duration (r = −0.37, p = 0.02; Fig. 2B), and showed a trend towards a negative correlation after controlling for age (partial correlation, r = −0.29, p = 0.06; Fig. 2C) as symptom duration and age correlated positively (r = 0.58, p = 0.001). The DLPFC GMV (age corrected) of FM patients did not correlate with any other characteristics (placebo effect [control-placebo]: intensity ratings, r = 0.14, p = 0.46, unpleasantness ratings, r = −0.08, p = 0.66; FIQ, r = 0.05, p = 0.82; daily pain, r = 0.01, p = 0.63. No other brain regions showed decreased or increased GMV in FM patients compared to controls.

**Figure 2.**
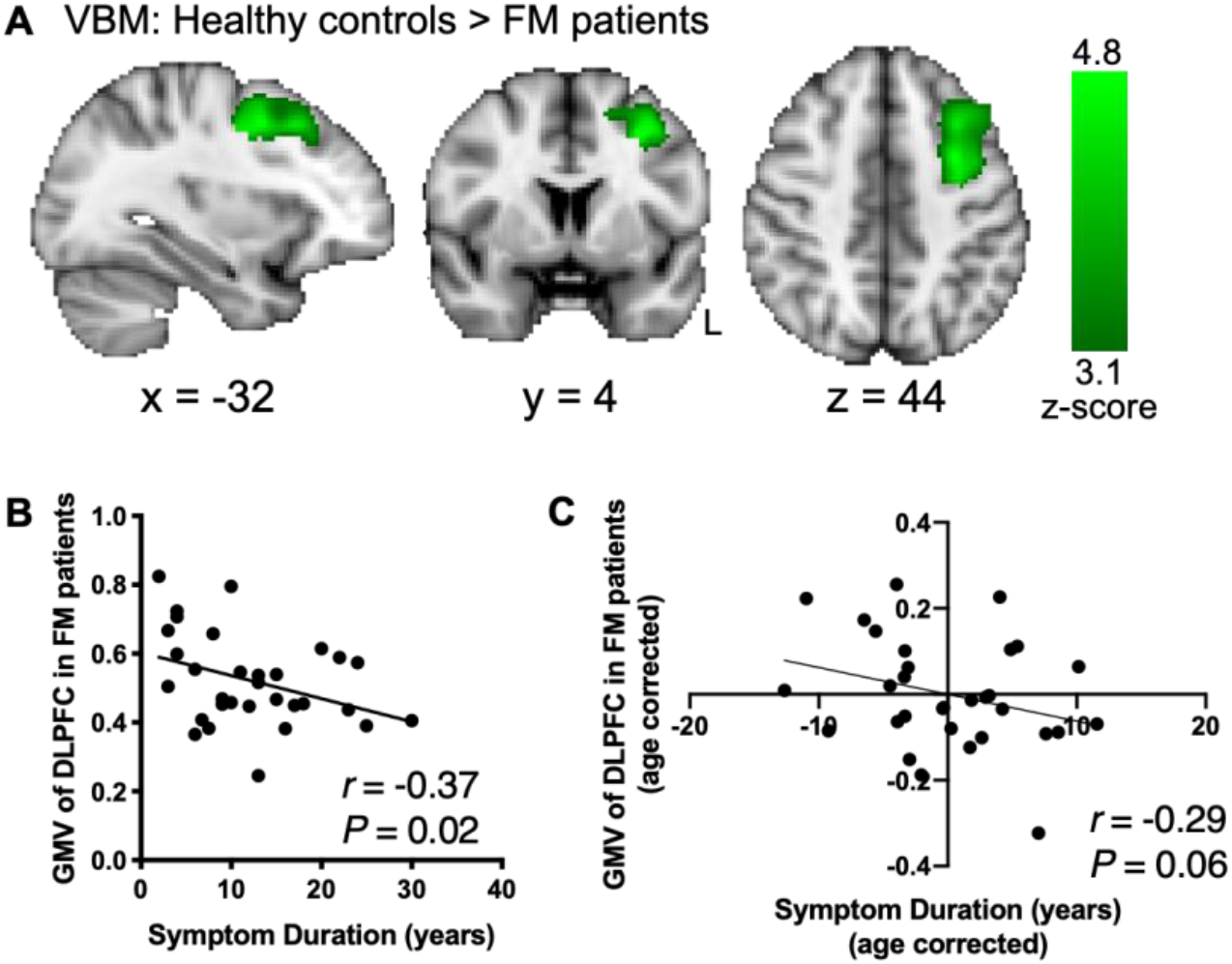
DLPFC GMV reduction in FM patients is related to symptom duration. A. VBM analysis of GMV comparing HCs and FM patients showed significant increases within the left dorsolateral prefrontal cortex (DLPFC) indicating DLPFC GMV reduction in FM patients (age corrected, cluster forming threshold z > 3.1, cluster correction p < 0.05; 1461 voxels; peak z-score = 4.8, p = 0.002; MNI coordinates −30, 4, 44). B. Average GMV of the DLPFC in FM patients significantly correlated negatively with the duration of fibromyalgia symptoms. C. A trends towards a negative correlation between DLPFC GMV and symptom duration remained after correcting for age. L, left.

### Q1: Magnitude of placebo analgesia did not differ between FM and HC groups

During the placebo test scans, participants received the same individualized heat temperatures for the control cream and placebo cream, a temperature midway between the individualized high and low temperatures presented during the conditioning scan. The temperatures did not differ between groups (HC 45.7°C ± 1.8°C, FM 46.1°C ± 1.38°C, p = 0.29).

A significant placebo analgesic effect was observed across all participants as indicated by reduced pain intensity ratings during the placebo cream condition compared to the control cream condition (Fig. 3A; placebo 135.6 ± 28.3, control 141.2 ± 25.1, F(1,76) = 6.03, p = 0.02, η_p2_ = 0.08). There was no main effect of group (HC 140.2 ± 29.1, FM 135.6 ± 23, F(1,76) = 0.6, p = 0.44, η_p2_ = 0.01) and no interactions were observed (cream x group, F(1,76) = 0.2, p = 0.66). More specifically, the difference between pain ratings during placebo and control did not differ between FM patients and HCs (Fig. 3B; control cream - placebo cream, HC 6.7 ± 18.9, FM 4.2 ± 19.9, p = 0.52; Cohen’s d = 0.146) indicating that the placebo effect in FM patients is comparable to that of healthy controls. A post-hoc power analysis revealed that with this effect size, 938 HC and 656 FM patients would be required to detect a significant cream x group interaction. Similarly, pain unpleasantness ratings significantly decreased during placebo across all participants (Fig. 3C; control 34.8 ± 22.2, placebo 28.9 ± 26.6, F(1,76) = 4.86, p = 0.03, η_p2_ = 0.07). A main effect of group was observed for pain unpleasantness, with FM patients rating pain across the placebo and control conditions as significantly less unpleasant than HCs (HC 36.6 ± 30.8, FM 25.6 ± 31.7, F(1,76) = 4.3, p = 0.04, η_p2_ = 0.06). However, no interaction between cream and group (F(1,76) = 0.58, p = 0.45, η_p2_ = 0.008) was observed indicating that the difference between pain unpleasantness ratings during placebo and control did not differ between FM patients and HCs (Fig. 3D; control cream - placebo cream, HC −7.9 ± 24, FM 3.4 ± 17.1, p = 0.42, Cohen’s d = 0.53).

**Figure 3.**
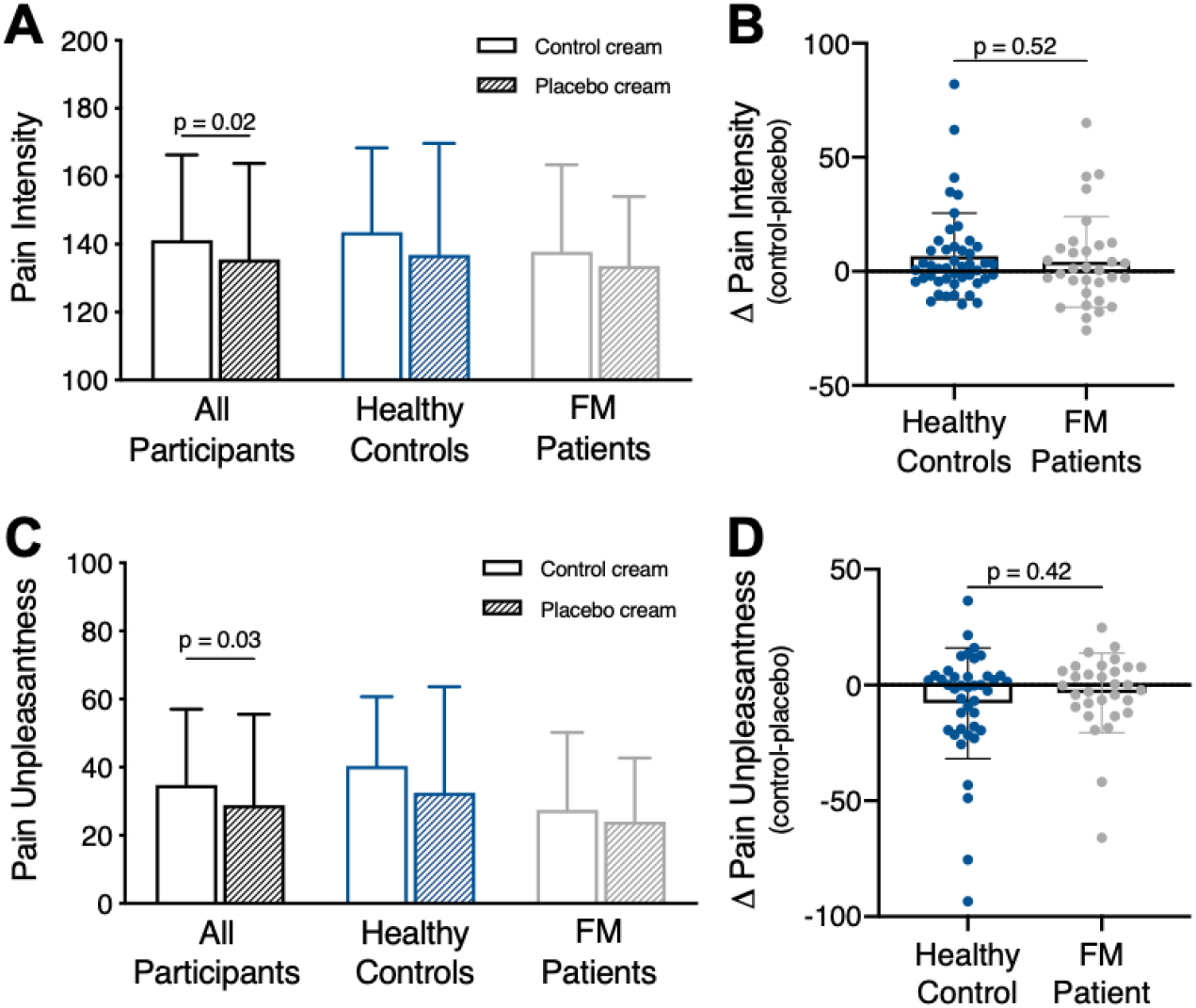
Behavioral placebo analgesic effects. Results are presented as mean±SD. A. Pain intensity ratings across all participants significantly decreased in response to the same temperature stimulus during the placebo cream condition compared to the control cream condition. Separate group plots show mean pain intensity ratings for each condition. B. Comparison of the pain intensity difference score (control cream - placebo cream) between HCs and FM patients shows no group differences in placebo effect. C. Pain unpleasantness ratings across all participants significantly decreased in response to the same temperature stimulus during the placebo cream condition compared to the control cream condition. Separate group plots show mean pain unpleasantness ratings for each condition. D. Group comparison of the pain unpleasantness difference score (control cream - placebo cream) shows no difference.

Consistent with ratings during the scan, when asked after the placebo test scan whether the placebo analgesic cream was effective, both groups reported slight to moderate effectiveness (HC 3.6 ± 2.9, FM 4.2 ± 2.3, p = 0.31, Cohen’s d = 0.23). The reported desire for pain relief was moderate and did not differ between groups during the placebo test scan (HC 5.3 ± 3.0, FM 5.2 ± 2.5, p = 0.79, Cohen’s d = 0.04).

### Q2: Brain regions involved in placebo analgesia did not differ between HC and FM

#### Pain-related activation

We examined pain-related activity across all participants during the placebo cream and control cream condition compared to baseline. Pain-related activations in the control cream condition showed typical patterns, including the ACC, and bilateral insula, S1, and S2 (z > 2.3, p < 0.05; Table 2; Fig. 4A). Similar regions were activated in the placebo cream condition (Fig. 4A). To determine whether placebo-related reductions in brain activity occurred, we examined the contrast “control cream > placebo cream” and found significantly more activation within the right mid-anterior and posterior insula (contralateral to the stimuli), and bilaterally in S2 during the control cream condition compared to placebo (Fig. 4B). The placebo-related reductions in the insula and S2 were comparable between the HC and FM groups (Fig. 4B inset). No increased activations were observed in pain modulatory regions or other brain regions (placebo cream > control cream; z > 2.3, p < 0.05). Similar to the behavioral results, there were no group differences or significant findings in the F-test for the cream*group interaction in neither the control cream > placebo cream contrast nor the placebo cream > control cream contrast (z > 2.3, p < 0.05; Table 2).

**Table 2.** Whole-brain BOLD responses during the first heat stimulation periods.

| Region | voxels | MNI |  |  | Peak<br>z-score | p-value |
| --- | --- | --- | --- | --- | --- | --- |
|  |  | x | y | z |  |  |
| **Control > baseline** |  |  |  |  |  |  |
| Insula (R) | 46636 | 28 | 65 | 39 | 9.01 | <0.001 |
| n. Accumbens (R) |  | 39 | 68 | 34 | 5.40 |  |
| ACC |  | 49 | 70 | 53 | 7.07 |  |
| Insula (L) |  | 60 | 72 | 38 | 8.05 |  |
| Occipital ctx (L) |  | 53 | 21 | 27 | 8.94 |  |
| Occipital ctx (R) |  | 28 | 17 | 43 | 8.33 |  |
| SI (R) |  | 37 | 43 | 70 | 6.84 |  |
| SII (L) |  | 74 | 52 | 44 | 7.77 |  |
| SII (R) |  | 18 | 51 | 45 | 7.74 |  |
| Thalamus (R) |  | 40 | 59 | 38 | 6.70 |  |
| **Placebo > baseline** |  |  |  |  |  |  |
| Planum polar | 43617 | 18 | 65 | 35 | 9.38 | <0.001 |
| n. Accumbens (R) |  | 40 | 67 | 33 | 6.54 |  |
| ACC |  | 49 | 73 | 51 | 7.16 |  |
| Insula (R) |  | 27 | 74 | 34 | 8.65 |  |
| Insula (L) |  | 64 | 54 | 39 | 7.15 |  |
| Occipital ctx (L) |  | 52 | 20 | 29 | 8.38 |  |
| Occipital ctx (R) |  | 31 | 16 | 43 | 7.68 |  |
| SI (R) |  | 39 | 44 | 70 | 6.15 |  |
| SII (L) |  | 76 | 51 | 44 | 6.50 |  |
| SII (R) |  | 15 | 49 | 47 | 6.81 |  |
| **Control > Placebo** |  |  |  |  |  |  |
| Occipital ctx (L) | 1710 | 53 | 12 | 37 | 4.69 | <0.001 |
| Occipital ctx (R) | 1624 | 37 | 14 | 39 | 5.89 | <0.001 |
| posterior Insula (R) | 1474 | 26 | 55 | 42 | 4.59 | <0.001 |
| mid-anterior Insula (R) |  | 27 | 70 | 38 | 3.74 |  |
| SII (R) |  | 20 | 49 | 45 | 4.17 |  |
| SII (L) | 1197 | 79 | 50 | 44 | 4.34 | <0.001 |
| **Placebo > Control** |  |  |  |  |  |  |
| - | - | - | - | - | - | - |
| **Group x cream (F-test)** |  |  |  |  |  |  |
| - | - | - | - | - | - | - |
| **Group x cream x drug (F-test)** |  |  |  |  |  |  |
| - | - | - | - | - | - | - |

**Figure 4.**
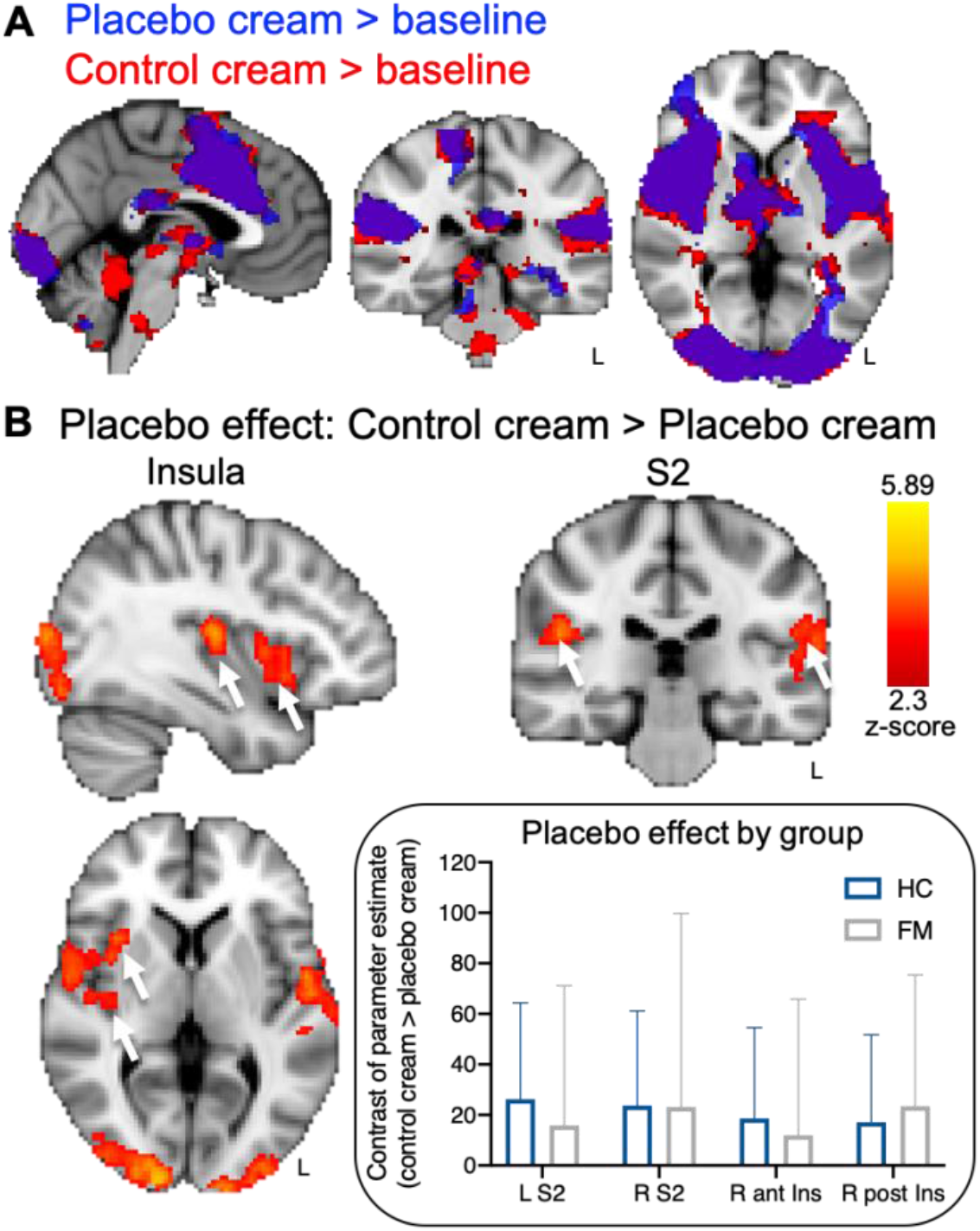
Neural placebo analgesic effects. A. Pain-related activations in response to the same temperature stimulus during the control cream condition (red) and placebo cream condition (blue) compared to baseline (overlapping regions displayed in purple). B. Significant placebo-related reductions (“placebo effect”, control > placebo) in brain activity were observed across all participants within the right mid-anterior and posterior insula and bilateral S2. Inset: The HC and FM patient contrast of parameter estimates for the placebo-modulated S2 and insula regions. All results are presented at a voxel-based threshold of z > 2.3, cluster correction of p < 0.05. ant, anterior; Ins, insula; L, left; post, posterior; R, right; S2, secondary somatosensory cortex.

#### NPS analysis

We then assessed whether the placebo condition modulated the NPS during the placebo test scan. We found a significantly smaller NPS response in the placebo cream condition compared to the control cream condition across all participants (control 1381.25 ± 945.08, placebo 997.83 ± 869.61, p < 0.001, Cohen’s d = 0.42). This finding corroborates the placebo analgesic effects on brain activation described in the previous paragraph. No group effect was observed (HC 531.4 ± 745.5, FM 170.8 ± 1139.2, p = 0.1, Cohen’s d = 0.4).

#### Anticipation-related activation

We examined activations during the anticipation periods (when subjects were looking at pictures of the cream containers) to uncover possible engagement of endogenous pain inhibiting brain networks during anticipation of pain relief. We found significant activation across all participants only in the occipital cortex during the anticipation period of the placebo cream condition compared to baseline (z > 2.3, p < 0.05; Fig. 5A) and compared to the anticipation of the control cream condition (z > 2.3, p < 0.05; Fig. 5B). No activation of pain modulatory regions previously described in the literature (e.g., DLPFC, VMPFC, ACC, PAG) was observed. During anticipation of the control cream condition compared to baseline (Fig. 5A), but not compared to anticipation of placebo, we found significant activation within the right hippocampus and right temporal cortex (z > 2.3, p < 0.05; Fig. 5A). No significant findings were observed in the F-test for the group*cream interaction for the anticipation of placebo cream > anticipation of control cream contrast or vice versa (z > 2.3, p < 0.05; Table 3).

**Figure 5.**
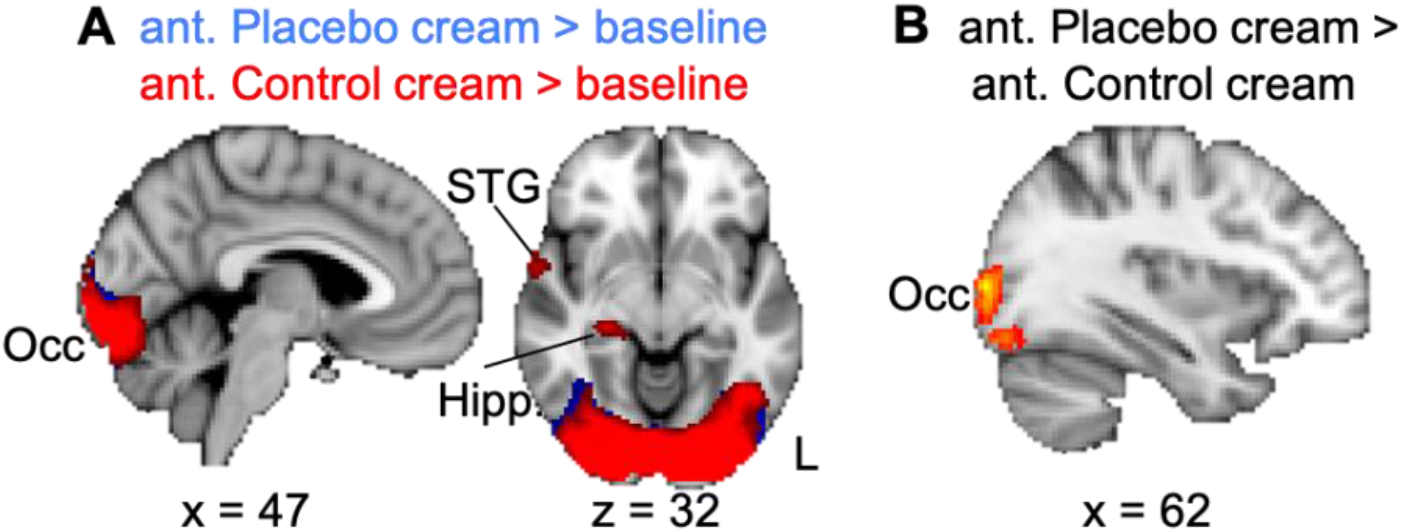
Anticipation-related activations. A. Compared to baseline, both anticipation conditions produced activation in the occipital cortex, and the STG and hippocampus were activated only during anticipation of the control cream condition. B. Greater activation of the occipital cortex was observed during anticipation of the placebo cream compared to anticipation of the control cream. No differences were observed in the inverse contrast. All results are presented at a voxel-based threshold of z > 2.3, cluster correction of p < 0.05. ant, anticipation; Hipp., hippocampus; L, left; STG; superior temporal gyrus.

**Table 3.** Whole-brain BOLD responses during the first anticipation periods.

| Region | voxels | MNI<br>coordinates |  |  | Peak<br>z-score | p-value |
| --- | --- | --- | --- | --- | --- | --- |
|  |  | x | y | z |  |  |
| **Control > baseline** |  |  |  |  |  |  |
| Occipital ctx (R) | 15608 | 39 | 14 | 38 | 9.73 | <0.001 |
| Hippocampus (R) |  | 32 | 50 | 32 | 4.25 |  |
| STG (R) |  | 15 | 63 | 36 | 4.13 | 0.038 |
| **Placebo > baseline** |  |  |  |  |  |  |
| Occipital ctx (L) | 16235 | 55 | 14 | 38 | 12.24 | <0.001 |
| Occipital fusiform ctx (R) |  | 34 | 17 | 32 | 9.85 |  |
| **Control > Placebo** |  |  |  |  |  |  |
| - | - | - | - | - | - | - |
| **Placebo > Control** |  |  |  |  |  |  |
| Occipital ctx (L) | 1229 | 62 | 17 | 39 | 4.17 | <0.001 |
| Occipital ctx (R) | 1056 | 32 | 17 | 38 | 4.14 | <0.001 |
| **Group x cream (F-test)** |  |  |  |  |  |  |
| - | - | - | - | - | - | - |
| **Group x cream x drug (F-test)** |  |  |  |  |  |  |
| - | - | - | - | - | - | - |

### Q3: Naloxone did not alter placebo analgesic responses

Naloxone did not alter the perceptual placebo effect compared to saline. There was no significant interaction for pain intensity ratings between the within-subject factor “cream” and the between-subject factor “drug” (cream*drug, F(1,76) = 0.08, p = 0.78, η_p2_ = 0.001). In addition, there was no interaction with group (group*drug, F(1,76) = 0.56, p = 0.46, η_p2_ = 0.01; cream*drug*group, F(1,76) = 0.26, p = 0.62, η_p2_ = 0.003). A post-hoc power analysis revealed that with these effect sizes, approximately 2.5 million participants would be required to reveal a significant cream*drug*group interaction. Similar results were found for pain unpleasantness (cream*drug, F(1,76) = 0.002, p = 0.96, η_p2_ = 0.000; group*drug, F(1,76) = 0.67, p = 0.67, η_p2_ = 0.003; cream*drug*group, F(1,76) = 0.41, p = 0.52, η_p2_ = 0.01).

Additionally, we found that naloxone compared to saline did not alter the neural placebo effect as no significant findings were observed in the F-test for the cream*drug or cream*drug*group interactions in the control cream > placebo cream contrast or vice versa (z > 2.3, p < 0.05; Table 2). Similarly, no drug effects were found on the anticipation of the placebo cream compared to the anticipation of the control cream for both interaction terms (z > 2.3, p < 0.05; Table 3).

#### NPS response

No significant NPS modulation by saline or naloxone was observed during placebo analgesia (control cream > placebo cream, Sal 402.43 ± 1030.36, Nal 367.96 ± 867.02, p = 0.87, Cohen’s d = 0.04).

## DISCUSSION

The present study investigated whether mechanisms of placebo analgesia in chronic pain patients with FM differed from healthy controls. Across all participants, we found reduced pain perception and pain-evoked functional activity within the right insula and bilateral secondary somatosensory cortex during placebo analgesia. Placebo modulation of nociceptive processing regions was further confirmed by a significant reduction in the NPS response. There were no differences between FM patients and healthy control participants in either pain intensity ratings or neural placebo-related effects. Across all conditions, FM patients reported the heat stimuli as less unpleasant than HCs, perhaps as a result of comparing the experimental pain to previous clinical pain episodes, as reported by over half of FM patients during a post-experimental interview session [38]. Lastly, there were no effects of naloxone administration nor any interactions between group and drug on anticipation-, placebo-, and NPS-related activations.

### Placebo effects on experimental pain in healthy and chronic pain populations

There was no difference in placebo analgesic responses in FM patients and HCs which is consistent with the few studies comparing placebo effects to experimental stimuli in pain patients and healthy participants. The most recent and largest behavioral study of this kind reported no differences in placebo effects between 363 chronic pain patients with temporomandibular disorder and 400 healthy controls [12]. Comparable placebo effects have also been reported between healthy participants and patients with episodic migraine [30], irritable bowel syndrome (IBS) [28], and atopic dermatitis [24]. Other studies have reported placebo effects to experimental stimuli in chronic pain patients without direct comparison to healthy participants [14; 33; 40], and a meta-analysis concluded that pain patients have a greater benefit from placebo treatment than healthy individuals [18]. These studies suggest that the presence of chronic pain might not alter placebo effects, despite the anatomical, neurochemical, and functional changes in the brain in chronic pain patients [3; 10; 11; 21; 22; 27; 31; 36; 47]. Nevertheless, evidence suggests that factors such as disease chronicity in patients with moderate to severe FM inversely correlates with placebo effects [25]. While we found no relationship between placebo effects and FM characteristics, future studies should compare cohorts with greater disease burden or pain severity to determine the extent to which these factors interact with placebo analgesia.

### Opioid versus non-opioid placebo analgesia

Several reports suggest that placebo analgesic responses are a result of accessing endogenous pain inhibitory opioid pathways [46; 49]. Here, the placebo effect was not blocked by the opioid antagonist, naloxone, suggesting that placebo analgesic effects in our study were opioid-independent. Although the dosage used in our study was lower than the dosage administered in two other studies with similar manipulations, i.e., verbal-suggestion and conditioning [1; 16], the dosage of approximately 10mg/kg is similar to that used in the first demonstration of naloxone reversal of placebo analgesia [29] and is a dose used clinically to reverse opioid-related overdoses. Further, we also concurrently examined affective touch perception in a sub-set (n = 52) of the participants and found that touch perception was altered by our naloxone dose [9], thus confirming that our dosage was physiologically effective. Some findings suggest that longer durations of experimental pain stimuli may allow for better engagement of opioid circuitry [6; 16; 39]. However, compared to Eippert et al. [16], we observed no naloxone effect despite having not only similar stimulus durations, but also a larger sample of healthy participants that received saline or naloxone. Another possibility that might account for the differences in results is the age and sex of the subjects. While our population was mainly middle-aged females, the Eippert population was exclusively young males.

An explanation for our lack of naloxone effects that we find the most compelling involves placebo induction via expectation versus conditioning. Naloxone has been shown to block placebo effects when placebo analgesia is created through suggestion-related expectations of pain relief but not when created through conditioning involving repeated experiences of pain relief under the same environmental contingencies [1; 40]. Unlike studies that just have explicit verbal suggestions telling participants that they are receiving a powerful pain-relieving analgesic [13; 16; 45], we added the additional instruction that naloxone might reverse the analgesic effects of the cream. Thus, the strong customary suggestion of impending pain relief may have been neutralized by the additional instruction. Nevertheless, our conditioning procedure involved the surreptitious reduction of the pain stimulus when paired with the placebo treatment. Many placebo analgesia studies incorporate this conditioning procedure along with high expectation-inducing verbal instructions [13; 16; 45]. However, our two days of conditioning, one of which occurred immediately prior to the experimental session, enhanced the conditioning component and simultaneously decreased the expectation component through our neutral verbal expectation instructions. While the inherent decrease in expectation could have diminished expectation-related components of the placebo effect [32], it is plausible that the repeated, multi-day conditioning-related placebo effects activated expectation-independent pathways. Thus, our observed opioid-independent placebo effects are likely influenced by learning-related conditioning that may not necessarily engage pain-relieving opioid pathways and, therefore, may neurobiologically differ from expectation-related responses.

Consistent with the idea that the current analgesic effects are not opioid-mediated is our finding that expectation-related circuitry was not significantly engaged in the present study. A meta-analysis of 25 neuroimaging studies of placebo analgesia and expectancy-based pain modulation identified regions consistently activated during expectation of pain relief, including the prefrontal cortex (DLPFC, VMPFC and orbitofrontal cortex), the midbrain surrounding the periaqueductal gray, and the rostral ACC [4]. Here, none of these regions were activated during the anticipation period preceding the stimulus on the “analgesic” skin.

### Does placebo analgesia involve bottom-up pain pathways?

A recent meta-analysis concluded that the mechanisms underlying placebo analgesia involve processes associated with the affective component of the pain experience, cognitive evaluation, pain-associated decision making, and mesolimbic reward processing, rather than engaging descending modulation onto afferent pain pathways [50]. This conclusion was based on the minimal placebo effects on the NPS within the 20 studies included in the meta-analysis and across all the studies using only placebo responders. This finding contradicts those reported by Eippert et al., in 2009 [17] where placebo analgesia decreased BOLD signals in the dorsal horn of the spinal cord, suggesting that placebo analgesia can modulate early nociceptive processes. Here, using the same multivariate brain activation pattern used in the meta-analysis [50], which includes brain regions involved in early nociceptive processing and is sensitive to intensities of evoked pain, we found that the NPS response [44] was greater during the control cream condition than the placebo cream condition, despite using the same temperature heat stimulus for both conditions and despite pain intensity ratings that differed by only ~5% between conditions. This suggests that placebo analgesia does, in part, alter bottom-up nociception, in contrast to the conclusion that “placebo treatments affect pain via brain mechanisms largely independent of effects on bottom-up nociceptive processing” [50]. Of the 20 studies included in the meta-analysis, 65% had a placebo-induction paradigm similar to ours, i.e., a combination of conditioning- and expectation-based induction, and 50% used a similar pain stimulus (heat pain), but our sample size was larger than 95% of the studies. Thus, it is possible that variations in study design (e.g. experimental pain type and duration, and placebo manipulations) and sample size could account for differences in NPS responses.

Additional caveats require discussion. First, due to protocol restrictions on subject replacements, we did not fully reach the objective of having 40 usable datasets in the FM group. Thus, given the subgroup sample sizes and limited number of heat pulses, it is possible that some analyses of interest may be underpowered which limits the generalizability of the results. Nevertheless, compared to similar studies with comparable or smaller sample sizes [16; 24; 28; 30; 40], no interactions showed any indication of a trend towards significance. Indeed, post-hoc power analyses indicated that we would need ~1600 participants to detect a group difference in placebo effect and ~2.5 million participants to detect a significant cream*group*drug interaction. Second, common verbal instructions used to induce high expectations of pain relief that engage prefrontal networks (e.g., DLPFC) and opioid pathways [19; 34; 43; 49] were likely neutralized by the additional instruction that naloxone may or may not block the “analgesic” effect of the cream. Thus, given the neural aberrancies in pathways overlapping with expectation-related placebo analgesia in FM patients [19; 43], e.g., decreased GMV within the DLPFC as observed in the FM patients in the present study, altered placebo effects under circumstances of high pain-relief expectations are still plausible.

In conclusion, the present findings provide evidence that opioid-independent predominantly conditioning-related placebo analgesia modulates pain perception and pain-evoked neural activity without accessing placebo pain modulatory networks in the prefrontal cortex. Chronic pain patients with FM did not differ from healthy participants in their behavioral or neural responses to the placebo manipulation, suggesting that the observed differences in prefrontal brain anatomy did not adversely affect conditioning-related placebo analgesia. This finding suggests that harnessing placebo effects based on conditioning in chronic pain patients might represent a relevant avenue for therapeutic strategies.

## Acknowledgements

This research was supported by the Intramural Research Program of the NIH, National Center for Complementary and Integrative Health. We thank Brian Walitt, Nicole Godwin, Linda Ellison-Dejewski, Brenda Justement, Susan Goo, and Patrick Korb for subject recruitment and clinical support, Lauren Atlas for guidance with the Neurological Pain Signature scripts, and Cortney Dable for assistance with the manuscript. The authors declare no conflicts of interest.

