## Supplementary Material for "Neural effects of placebo analgesia in fibromyalgia patients and healthy individuals"

1. Reported side effects of naloxone and saline, Table 1.
2. Correlations between FM characteristics and pain ratings, Table 2.
3. Results of heat pulse 2 of first placebo test scan, Figure 1, Table 3.
4. Results of second placebo test scan, Table 4, Table 5.

#### 1. Reported side effects of naloxone and saline.

Participants reported few to no side effects from either drug. Results are presented as mean (SD). The scale used to rate adverse effects of the drugs ranged from 0 (non-existent) to 6 (extremely strong).

Table 1. Reported side effects of naloxone and saline.

| | FM (n=32) | | HC (n=46) | |
|---|---|---|---|---|
| | saline | naloxone | saline | naloxone |
| Blurred vision | 0 | 0 | 0 | 0.2(1.04) |
| Dizziness | 0 | 0 (0.3) | 0 | 0 |
| Dry mouth | 0.2(0.7) | 0.7(1.5) | 0.1(0.4) | 0.3(0.9) |
| Dry skin | 0.1(0.3) | 0.1(0.2) | 0 | 0 |
| Headache | 0.2(0.6) | 0.4(1.4) | 0.1(0.5) | 0 |
| Nausea | 0 | 0 | 0 | 0(0.2) |
| Sedation | 0 | 0.4(0.9) | 0.1(0.4) | 0.6(1.3) |

### 2. Correlations between FM characteristics and pain ratings

Considering the evidence indicating that FM duration decreases placebo effects [22], we assessed the FM characteristics below to determine whether they related to FM placebo analgesic effects. We found no evidence of trends in the present FM cohort indicative of placebo response changing as a function of FM characteristics. As expected, the only significant positive correlation was found between placebo pain intensity and unpleasantness ratings.

Table 2. Pearson correlation matrix of FM characteristics and placebo response (pain ratings: control-placebo).

| | | M | SD | 1 | 2 | 3 | 4 | 5 |
|---|---|---|---|---|---|---|---|---|
| 1 | *Placebo response (intensity) | 4.15 | 19.87 | -- | | | | |
| 2 | ◊Placebo response (unpleasantness) | 24.01 | 18.67 | 0.77 (<0.001) | -- | | | |
| 3 | Symptom duration | 11.91 | 7.48 | 0.19 (0.30) | 0.19 (0.30) | -- | | |
| 4 | FIQ | 41.97 | 19.32 | 0.30 (0.13) | 0.15 (0.48) | -0.13 (0.51) | -- | |
| 5 | Daily pain | 6.63 | 2.52 | 0.02 (0.89) | 0.13 (0.52) | -0.20 (0.33) | 0.41 (0.06) | -- |

Results are presented as r(*p*).

*Difference between control pain intensity rating and placebo.

◊Difference between control pain unpleasantness rating and placebo.

Symptom duration, N=32

FIQ, Fibromyalgia Impact Questionnaire, N=26

Daily pain, N=26

#### 3. Results of heat pulse 2 of first placebo test scan

The neural results of the second anticipation period and heat pulse (black arrows in Fig. 1) of the first placebo test scan are summarized below in Table 3.

*Heat stimulation period.* As expected, the heat stimulation presented during the placebo and control cream conditions each produced significant activations in pain-related regions (e.g., insula, ACC, SII) when compared to baseline. However, as indicated by the control cream > placebo cream contrast, no placebo-related decreases were observed in any brain regions during heat pulse 2 of each trial. Placebo-related increases (i.e., placebo cream > control cream) were observed only in the frontal pole. No group or drug differences were found, and no group * cream interactions were observed. While we did observe a significant group * cream * drug interaction in the control cream > placebo cream contrast, we found that the interaction was produced when HCs with saline were grouped with FM patients with naloxone and then compared to HCs with naloxone grouped with FM patients with saline, i.e., $HC_{sal} + FM_{nal} > HC_{nal} + FM_{sal}$. For the purposes of the present study, we concluded that this was not scientifically meaningful.

*Anticipation period.* Similarly, the second anticipation period preceding heat pulse 2 produced activation within pain-related regions and also within frontal regions in the control cream and placebo cream conditions compared to baseline. However, only differences in within the occipital pole and intracalcarine cortex were observed in the control cream > placebo cream contrast. No expectation-related activations or any other activations were observed in the placebo cream > control cream contrast, and no interactions were observed.

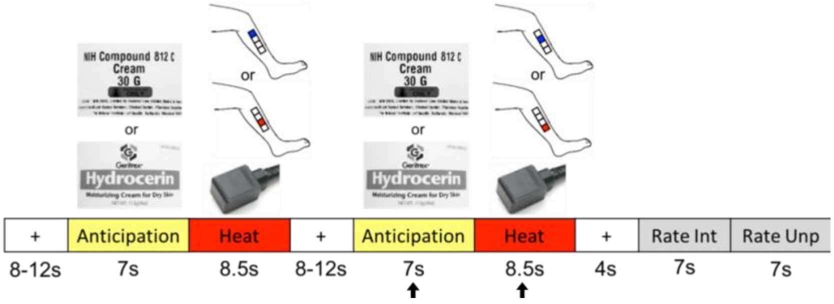

**Figure 1.** Trial paradigm for experimental scan. Black arrows indicate second heat pulse.

**Table 3.** Whole-brain BOLD responses during heat pulse 2 of the first placebo experimental scan.

| Region | voxels | MNI coordinates x | y | z | Peak z-score | p-value |
|---|---|---|---|---|---|---|
| ***Anticipation period*** | | | | | | |
| **Control > baseline** | | | | | | |
| Occipital pole (L) | 48025 | 52 | 12 | 37 | 11.80 | <0.001 |
| Hippocampus (L) | | 56 | 48 | 32 | 6.39 | |

| | | | | | | |
|---|---|---|---|---|---|---|
| Inferior frontal gyrus (L) | | 67 | 66 | 46 | 4.49 | |
| post. Insula (R) | | 27 | 54 | 42 | 5.58 | |
| Frontal medial ctx (L) | | 49 | 82 | 27 | 5.02 | |
| Occipital pole (R) | | 36 | 19 | 32 | 11.50 | |
| SII (L) | | 76 | 51 | 44 | 6.18 | |
| SII (R) | | 18 | 52 | 45 | 5.67 | |
| Thalamus (L) | | 50 | 53 | 38 | 3.79 | |
| Thalamus (R) | | 38 | 51 | 37 | 3.91 | |
| SI (R) | 490 | 38 | 41 | 69 | 4.59 | 0.01 |
| **Placebo > baseline** | | | | | | |
| Occipital pole (L) | 39683 | 59 | 18 | 30 | 9.95 | <0.001 |
| Hippocampus (L) | | 56 | 49 | 32 | 5.70 | |
| mid Insula (L) | | 64 | 61 | 41 | 3.26 | |
| Paracingulate gyrus (R) | | 39 | 86 | 47 | 3.01 | |
| Occipital pole (R) | | 41 | 18 | 32 | 9.53 | |
| SII (R) | | 20 | 50 | 44 | 5.88 | |
| SII (L) | | 76 | 49 | 44 | 4.48 | |
| Thalamus (L) | | 49 | 54 | 38 | 3.68 | |
| Frontal pole (L) | 439 | 51 | 94 | 47 | 3.73 | 0.029 |
| Frontal pole (R) | | 41 | 93 | 43 | 3.28 | |
| **Control > Placebo** | | | | | | |
| Intracalcarine ctx (R) | 616 | 37 | 19 | 40 | 3.96 | 0.002 |
| Occipital pole (R) | | 36 | 16 | 46 | 3.64 | |
| **Placebo > Control** | | | | | | |
| - | - | - | - | - | - | - |
| **Group x cream (F-test)** | | | | | | |
| - | - | - | - | - | - | - |
| **Group x cream x drug (F-test)** | | | | | | |
| - | - | - | - | - | - | - |
| ***Stimulation period*** | | | | | | |
| **Control > baseline** | | | | | | |
| Temporal pole (R) | 66991 | 16 | 68 | 32 | 9.61 | <0.001 |
| ACC | | 45 | 67 | 57 | 7.91 | |
| Central operc. ctx (R) | | 21 | 66 | 37 | 8.57 | |
| post. Insula (L) | | 63 | 55 | 36 | 7.63 | |
| ant. Insula (R) | | 28 | 72 | 37 | 9.53 | |
| post. Insula (R) | | 28 | 53 | 40 | 8.84 | |
| Precentral gyrus (R) | | 16 | 67 | 37 | 8.64 | |
| Occipital ctx (L) | | 60 | 18 | 41 | 8.45 | |
| Occipital ctx (R) | | 39 | 18 | 33 | 8.60 | |
| SI (R) | | 36 | 44 | 70 | 7.08 | |
| SII (L) | | 72 | 53 | 43 | 7.95 | |
| SII (R) | | 15 | 53 | 44 | 8.08 | |

| | | | | | | |
|---|---|---|---|---|---|---|
| Thalamus (R) | | 39 | 57 | 36 | 6.51 | |
| **Placebo > baseline** | | | | | | |
| Occipital ctx (R) | 77149 | 40 | 15 | 38 | 10.50 | <0.001 |
| ACC | | 46 | 67 | 57 | 9.51 | |
| Central operc. ctx (R) | | 17 | 66 | 36 | 9.68 | |
| ant. Insula (L) | | 64 | 66 | 37 | 8.22 | |
| ant. Insula (R) | | 28 | 74 | 34 | 9.55 | |
| Occipital ctx (L) | | 48 | 16 | 32 | 8.85 | |
| SI (R) | | 36 | 44 | 69 | 7.50 | |
| SII (L) | | 74 | 50 | 44 | 7.63 | |
| SII (R) | | 15 | 51 | 46 | 8.75 | |
| SMG (R) | | 13 | 50 | 47 | 9.93 | |
| **Control > Placebo** | | | | | | |
| - | - | - | - | - | - | - |
| **Placebo > Control** | | | | | | |
| Frontal pole (R) | 791 | 25 | 85 | 43 | 4.06 | <0.001 |
| **Group x cream (F-test)** | | | | | | |
| - | - | - | - | - | - | - |
| **Group x cream x drug (F-test)** | | | | | | |
| Precuneous ctx | 2912 | 45 | 34 | 56 | 4.34 | <0.001 |
| Lat. Occipital ctx (R) | | 20 | 29 | 48 | 4.04 | |
| Lat. Occipital ctx (L) | 1774 | 65 | 28 | 53 | 4.47 | <0.001 |
| Angular gyrus (L) | | 70 | 37 | 49 | 3.95 | |
| Supramarginal gyrus (L) | | 70 | 40 | 59 | 3.96 | |
| Middle frontal gyrus (L) | 1067 | 65 | 67 | 52 | 4.06 | <0.001 |
| Superior frontal gyrus | | 54 | 78 | 58 | 3.78 | |

"-", no significant results; ACC, anterior cingulate cortex; ant., anterior; ctx., cortex; L, left; Lat., lateral; post., posterior; R, right; operc., operculum; SI, primary somatosensory cortex; SII, secondary somatosensory cortex; SMG; supramarginal gyrus.

### 4. Results of second placebo test scan

*Behavioral results.* For the second placebo experimental scan, the cream treated regions of the left leg (proximal vs. distal) were counterbalanced with the first placebo experimental scan. The participants received the same individualized heat temperatures as the first experimental scan. Although participants rated pain lower when the heat stimulus was presented to the placebo cream site than the control site, there was no significant main effect of cream (pain intensity: control 129.5±4.2, placebo 125.3±3.5, p=0.15). Furthermore, there was no difference between FM and HC participants (pain intensity: HC 128±4.5, FM 126.8±5.6, p=0.87), and no effect of naloxone (pain intensity: Sal 125.9±5.4, Nal 128.8±4.7, p=0.69). Finally, no significant interactions were found (pain intensity: cream*group, p=0.68; cream*drug, p=0.64; group*drug, p=0.8; cream*group*drug, p=0.1). Similarly, we found no significant effects on pain unpleasantness (cream: control -23.7±3.4, placebo -19.6±3.3, p=0.15; group: HC -26.7±4, FM -16.6±4.7, p=0.11; drug: Sal -19±4.7, Nal -24.3±4, p=0.4; cream*group, p=0.85; cream*drug, p=0.27 group*drug, p=0.83; cream*group*drug, p=0.14).

<u>Neural response to heat pulse 1 of second test scan</u>

*Heat stimulation period.* As summarized in Table 4, the heat stimuli presented during both the control cream and placebo cream conditions produced pain-related activations (e.g., insula and SII) compared to baseline. In the control cream > placebo cream contrast during stimulation, which would indicate placebo-related reductions, we did not observe differences in any of these pain-related regions. We did observe significantly greater activations within the lingual gyrus and occipital cortex, which might relate to differences in visualization. Whereas in the first scan we did not observe any regions with significantly greater activation during the placebo condition than during the control condition, during the second scan, we found significant differences in several brain regions, including S1, despite lower pain ratings during placebo. No differences for group or drug were found, and no interactions were observed.

*Anticipation period.* Differences in the placebo cream > control cream contrast were only observed within the occipital cortex during the anticipation condition. No differences for group or drug were found, and no interactions were observed.

<u>Neural response to heat pulse 2 of second test scan</u>

*Heat stimulation period.* Table 5 shows pain-related activations in response to the heat stimuli during the control and placebo cream condition, similar to the activation patterns observed during the other heat pulses. However, no placebo-related reductions were observed in any brain regions (control > placebo contrast), and placebo-related increases (placebo > control) were only observed in the lingual gyrus. No differences for group or drug were found, and no interactions were observed.

*Anticipation period.* Significant differences were observed only within the lateral occipital cortex in the placebo cream > control cream contrast. Similar to the first anticipation period and the anticipation periods of the first scan, we did not observe any differences in placebo expectation-related regions. No differences for group or drug were found, and no interactions were observed.

Habituation to the heat stimuli may account for the lack of differences observed during the second experimental run, as pain intensity ratings across all participants and conditions (group, drug, and cream) significantly decreased between the first and second placebo scans (pain intensity ratings across groups and conditions, placebo scan 1, 137.9±3, placebo scan 2, 127.4±3.6, p<0.001).

**Table 4.** Whole-brain BOLD responses of the first anticipation and heat pulse of the second placebo experimental scan.

| Region | voxels | MNI coordinates | | | Peak z-score | p-value |
|---|---|---|---|---|---|---|
| | | x | y | z | | |
| ***Anticipation period*** | | | | | | |
| **Control > baseline** | | | | | | |
| Occipital ctx (L) | 14716 | 47 | 18 | 29 | 12.2 | <0.001 |
| Occipital ctx (R) | | 38 | 15 | 35 | 11 | |
| SII (R) | 412 | 21 | 49 | 46 | 4.48 | 0.039 |
| **Placebo > baseline** | | | | | | |
| Frontal pole (L) | 549 | 49 | 91 | 45 | 4.16 | 0.008 |
| Frontal pole (R) | | 39 | 95 | 40 | 3.75 | |
| Occipital ctx (L) | 28953 | 52 | 17 | 28 | 12.8 | <0.001 |
| Occipital ctx (R) | | 40 | 14 | 42 | 10.24 | |
| SII (R) | 1290 | 23 | 48 | 45 | 5.22 | <0.001 |
| SI (R) | 436 | 34 | 43 | 69 | 4.57 | 0.029 |
| Subcallosal ctx | 2385 | 42 | 71 | 31 | 4.38 | <0.001 |
| Frontal medial ctx | | 49 | 86 | 28 | 4.20 | |
| STG (R) | 1852 | 14 | 64 | 36 | 4.80 | <0.001 |
| MTG (R) | | 14 | 60 | 29 | 4.23 | |
| **Control > Placebo** | | | | | | |
| - | - | - | - | - | - | - |
| **Placebo > Control** | | | | | | |
| Lateral occipital ctx (R) | 2766 | 26 | 20 | 42 | 4.72 | <0.001 |
| Occipital fusiform ctx (L) | 2890 | 65 | 27 | 30 | 5.17 | <0.001 |
| **Group x cream (F-test)** | | | | | | |
| - | - | - | - | - | - | - |
| **Group x cream x drug (F-test)** | | | | | | |
| - | - | - | - | - | - | - |
| ***Stimulation period*** | | | | | | |
| **Control > baseline** | | | | | | |
| Occipital ctx (R) | 9389 | 38 | 14 | 42 | 8.99 | <0.001 |
| Lingual gyrus (L) | | 48 | 20 | 28 | 8.79 | |
| Occipital fusiform gyrus (L) | | 59 | 22 | 28 | 8.55 | |
| Paracingulate (R) | 4189 | 44 | 71 | 56 | 6.72 | <0.001 |
| ACC (R) | | 43 | 70 | 52 | 5.87 | |
| SI (R) | | 35 | 44 | 70 | 6.03 | |
| STG (R) | 6574 | 15 | 63 | 35 | 8.20 | <0.001 |
| Insula (anterior) (R) | | 25 | 71 | 32 | 8.03 | |
| Insula (posterior) (R) | | 28 | 55 | 42 | 7.60 | |
| SII (R) | | 17 | 52 | 47 | 6.77 | |

| | | | | | | |
|---|---|---|---|---|---|---|
| STG (L) | 5449 | 75 | 62 | 35 | 7.48 | <0.001 |
| SII (L) | | 74 | 49 | 44 | 7.39 | |
| Planum polar (L) | | 72 | 63 | 35 | 7.16 | |
| Insula (ant) (L) | | 61 | 72 | 37 | 7.13 | |
| Thalamus (R) | 1015 | 40 | 57 | 39 | 5.01 | <0.001 |
| Thalamus (L) | | 48 | 58 | 35 | 3.86 | |
| **Placebo > baseline** | | | | | | |
| Insula (L) | 5523 | 61 | 74 | 37 | 6.34 | <0.001 |
| Occipital ctx (L) | 16245 | 55 | 23 | 28 | 9.30 | <0.001 |
| Occipital ctx (R) | | 34 | 14 | 44 | 8.42 | |
| Planum Polar (R) | 13921 | 16 | 62 | 36 | 8.41 | <0.001 |
| Central opercular ctx (R) | | 19 | 63 | 38 | 7.76 | |
| Insula (anterior) (R) | | 26 | 67 | 40 | 7.43 | |
| Insula (posterior) (R) | | 27 | 54 | 38 | 7.11 | |
| Paracingulate/ACC | | 43 | 72 | 55 | 7.20 | |
| SI (R) | | 35 | 43 | 70 | 6.19 | |
| SII (R) | | 21 | 48 | 47 | 6.60 | |
| Thalamus (R) | | 39 | 57 | 35 | 4.69 | |
| **Control > Placebo** | | | | | | |
| Lingual gyrus (L) | 467 | 51 | 20 | 30 | 4.57 | 0.018 |
| Occipital ctx (R) | 434 | 35 | 14 | 40 | 4.87 | 0.027 |
| **Placebo > Control** | | | | | | |
| Cerebellum (R) | 566 | 43 | 23 | 23 | 3.42 | 0.006 |
| Lateral occipital ctx (R) | 637 | 14 | 32 | 33 | 3.45 | 0.003 |
| MTG (R) | | 13 | 48 | 31 | 3.40 | |
| SPL (R) | 1148 | 30 | 36 | 65 | 4.23 | <0.001 |
| SI (R) | | 24 | 48 | 57 | 3.94 | |
| SMG (R) | | 24 | 42 | 59 | 3.53 | |
| **Group x cream (F-test)** | | | | | | |
| - | - | - | - | - | - | - |
| **Group x cream x drug (F-test)** | | | | | | |
| - | - | - | - | - | - | - |

"-", no significant results; (L), left; (R), right; ACC, anterior cingulate cortex; ctx, cortex; MTG; middle temporal gyrus; SI, primary somatosensory cortex; SII, secondary somatosensory cortex; n., nucleus; SMG; supramarginal gyrus; SPL, superior parietal lobe; STG, superior temporal gyrus.

**Table 5.** Whole-brain BOLD responses during heat pulse 2 of the second placebo experimental scan.

| Region | voxels | MNI coordinates | | | Peak z-score | p-value |
|---|---|---|---|---|---|---|
| | | x | y | z | | |
| ***Anticipation period*** | | | | | | |
| **Control > baseline** | | | | | | |
| Occipital fusiform gyrus (R) | 27121 | 36 | 18 | 27 | 11.16 | <0.001 |
| Frontal medial ctx | | 48 | 80 | 25 | 4.52 | |
| Occipital pole (L) | | 55 | 14 | 35 | 10.40 | |
| Occipital pole (R) | | 37 | 13 | 37 | 9.54 | |
| SI (L) | 481 | 77 | 46 | 46 | 3.91 | 0.019 |
| Heschl's gyrus | | 73 | 55 | 40 | 3.15 | |
| STG (L) | | 75 | 63 | 35 | 3.43 | |
| Temporal pole (L) | | 74 | 66 | 34 | 3.87 | |
| Precentral gyrus (L) | 459 | 75 | 64 | 52 | 4.92 | 0.025 |
| **Placebo > baseline** | | | | | | |
| Occipital pole (R) | 43957 | 40 | 16 | 34 | 13.45 | <0.001 |
| Frontal medial ctx | | 49 | 82 | 27 | 3.89 | |
| Lingual gyrus (L) | | 47 | 19 | 28 | 11.31 | |
| Precentral gyrus | | 71 | 65 | 56 | 4.66 | |
| SI (R) | | 39 | 41 | 70 | 3.66 | |
| SII (L) | | 72 | 49 | 43 | 4.24 | |
| SII (R) | | 22 | 48 | 47 | 5.30 | |
| ACC | 462 | 45 | 59 | 59 | 3.75 | 0.013 |
| **Control > Placebo** | | | | | | |
| - | - | - | - | - | - | - |
| **Placebo > Control** | | | | | | |
| Lat. occipital ctx (L) | 444 | 59 | 18 | 27 | 3.40 | 0.013 |
| **Group x cream (F-test)** | | | | | | |
| - | - | - | - | - | - | - |
| **Group x cream x drug (F-test)** | | | | | | |
| - | - | - | - | - | - | - |
| ***Stimulation period*** | | | | | | |
| **Control > baseline** | | | | | | |
| ant. Insula (R) | 38407 | 26 | 68 | 39 | 8.01 | <0.001 |
| post. Insula (L) | | 64 | 54 | 38 | 7.56 | |
| Central operc. ctx (R) | | 19 | 66 | 38 | 7.89 | |
| Frontal pole (R) | | 25 | 84 | 49 | 4.82 | |
| Occipital pole (R) | | 34 | 17 | 36 | 7.98 | |

| | | | | | | |
|---|---|---|---|---|---|---|
| Planum polar (L) | | 70 | 62 | 36 | 7.94 | |
| SII (L) | | 75 | 50 | 45 | 6.65 | |
| SII (R) | | 20 | 48 | 46 | 6.96 | |
| Thalamus (L) | | 51 | 55 | 39 | 4.69 | |
| Thalamus (R) | | 39 | 58 | 38 | 5.88 | |
| Paracingulate gyrus | 5416 | 42 | 70 | 58 | 8.72 | <0.001 |
| ACC | | 43 | 72 | 52 | 6.03 | |
| SI (R) | | 35 | 44 | 70 | 6.16 | |
| **Placebo > baseline** | | | | | | |
| Lingual gyrus (L) | 71439 | 47 | 18 | 32 | 10.59 | <0.001 |
| ACC | | 43 | 69 | 53 | 6.42 | |
| Frontal pole (R) | | 22 | 81 | 47 | 6.54 | |
| ant. Insula (R) | | 27 | 70 | 39 | 8.96 | |
| ant. Insula (L) | | 62 | 70 | 38 | 7.67 | |
| Occipital pole | | 45 | 17 | 32 | 10.08 | |
| Paracingulate gyrus | | 44 | 70 | 58 | 8.06 | |
| SI (R) | | 37 | 45 | 71 | 6.09 | |
| SII (L) | | 75 | 49 | 46 | 7.01 | |
| SII (R) | | 21 | 48 | 47 | 7.34 | |
| Temporal pole (R) | | 16 | 68 | 32 | 9.12 | |
| Thalamus (L) | | 49 | 55 | 34 | 5.65 | |
| Thalamus (R) | | 40 | 54 | 34 | 6.01 | |
| SI (L) | 487 | 55 | 40 | 69 | 4.66 | 0.031 |
| **Control > Placebo** | | | | | | |
| - | - | - | - | - | - | - |
| **Placebo > Control** | | | | | | |
| Lingual gyrus (R) | 743 | 42 | 28 | 30 | 4.10 | <0.001 |
| **Group x cream (F-test)** | | | | | | |
| - | - | - | - | - | - | - |
| **Group x cream x drug (F-test)** | | | | | | |
| - | - | - | - | - | - | - |

"-", no significant results; ACC, anterior cingulate cortex; ant., anterior; ctx., cortex; L, left; Lat., lateral; post., posterior; R, right; operc., operculum; SI, primary somatosensory cortex; SII, secondary somatosensory cortex; STG, superior temporal gyrus.
